# Tapping into the native *Pseudomonas* Bacterial Biofilm Structure by High-Resolution 1D and 2D MAS solid-state NMR

**DOI:** 10.1101/2023.10.02.560490

**Authors:** Chang-Hyeock Byeon, Ted Kinney, Hakan Saricayir, Sadhana Srinivasa, Meghan K. Wells, Wook Kim, Ümit Akbey

## Abstract

We present a high-resolution 1D and 2D magic-angle spinning (MAS) solid-state NMR (ssNMR) study to characterize native *Pseudomonas fluorescens* colony biofilms at natural abundance without isotope-labelling. By using a high-resolution INEPT-based 2D ^1^H-^13^C ssNMR spectrum and thorough peak deconvolution approach at the 1D ssNMR spectra, approximately 80/134 (in 1D/2D) distinct biofilm chemical sites were identified. We compared CP and INEPT ^13^C ssNMR spectra to different signals originating from the mobile and rigid fractions of the biofilm, and qualitative determined dynamical changes by comparing CP buildup behaviors. Protein and polysaccharide signals were differentiated and identified by utilizing FapC signals as a template, a biofilm forming functional amyloid from *Pseudomonas*. We also attempted to identify biofilm polysaccharide species by using ^1^H/^13^C chemical shifts obtained from the 2D spectrum. This study marks the first demonstration of high-resolution 2D ssNMR spectroscopy for characterizing native bacterial biofilms and expands the scope of ssNMR in studying biofilms. Our experimental pipeline can be readily applied to other in vitro biofilm model systems and natural biofilms and holds the promise of making a substantial impact on biofilm research, fostering new ideas and breakthroughs to aid in the development of strategic approaches to combat infections caused by biofilm-forming bacteria.

## 1. Introduction

Many species of bacteria form densely structured communities known as biofilms that confer protection to the resident cells against diverse environmental stressors. Biofilm protected bacteria cause ∼80% of all chronic infections, which are difficult to treat and augment antimicrobial-resistance (AMR).^1-3^ AMR results in ∼one million casualties per year and is estimated to cause more death than cancer by 2050.^4^ The structural integrity of biofilms is maintained by a complex array of extracellular polymeric compounds,^5^ including proteins in the form of amyloid fibrils and polysaccharides, but very little structural information exists about biofilms and their components. For the protein part of biofilms, currently, there is no high-resolution structure of a biofilm forming functional amyloid other than TasA from *Bacillus* determined from *in vitro* or purified preparations.^6, 7^ Similarly, only limited structural information exists on the polysaccharide fraction of biofilms, which is a complex environment for *Pseudomonas* species composed of Psl, Pel, alginate, and other polysaccharides.^8, 9^ This lack of structural understanding of biofilms, hampers development of therapeutic strategies. Novel approaches directed by structural insights are necessary for obtaining detailed structural information on biofilms and structure-activity relationship, that will be instrumental in advancing the fight against chronic infections and AMR.

Structural biology has celebrated many achievements in recent years, but studying proteins under native physiological conditions, such as in biofilms, remains a significant challenge. High-resolution techniques mostly rely on unnaturally high concentrations and quantities of protein. For characterization of biological systems at their native environment, magic-angle spinning (MAS) solid-state NMR (ssNMR) and cryo-EM/tomography are very powerful tools, and could provide atomic or near-atomic resolution information, respectively.^10-14^ Compared to other techniques, ssNMR is extremely powerful if/when the sensitivity problem is addressed. In particular, proton-detected (^1^H-detected) ssNMR and DNP-enhanced ssNMR (DNP ssNMR here on) are potentially game-changing in studying protein structure in native biofilms due to the remarkable increase in ssNMR sensitivity that reduces sample requirements by several orders of magnitude compared to conventional ssNMR.^15-17^ Methodological developments in sample preparation/optimization and technological breakthroughs in DNP-enhanced ssNMR have been demonstrated on complex systems in native environments, whole cells, and extracts.^18-20^ We provided the first example of measuring the deuterated functional amyloid TasA in its native *Bacillus* bacterial biofilm and compared to *in vitro* preparations by using proton-detected high-resolution ssNMR.^21^

Over the past decade, ssNMR spectroscopy has been widely used to characterize bacterial, plant, and fungal cell walls.^22-24^ However, particularly in biofilm research, ssNMR studies thus far relied primarily on 1D ssNMR spectroscopy, which often poses challenges such as signal overlap and limited quantification due to lower-resolution and signal overlap. Nevertheless, structural and compositional information could be obtained and correlated to the nature of bacterial and fungal biofilms.^22, 23, 25^ To overcome the limitations due to 1D ssNMR spectroscopy and differentiate signals from different biofilm parts, e.g. from proteins and polysaccharides, nD ssNMR experiments is crucial and proven powerful in plant and fungal cell wall studies.^26, 27^ To the best of our knowledge, there are only a few multidimensional ssNMR examples applied; by conventional ssNMR on cell-walls of *S. aureus*, high-resolution fast-MAS on *E. coli*-*P. aureginosa* cell-wall, and high-sensitivity DNP-enhanced ssNMR on *B. subtilis* cell-wall.^28-30^ The ^13^C-^13^C DARR ssNMR was used to observe NMR spectrum on a fully labeled *S. aureus* cell walls, and tentative assignments was used to understand structural details.^29^ The DNP enhanced ssNMR experiments have been demonstrated utilizing 2D spectra on native systems like fungi and plants.^24^ This high-sensitivity DNP approach has shown great utility in characterizing, e.g., intact or extracted bacterial cell walls as a sole demonstration till now.^28^ Similarly, there are only few examples of utilizing high-resolution proton and carbon-detected ssNMR for bacterial cell-wall characterization.^30, 31^ In summary, a small number of multidimensional ssNMR results highlights that there is a knowledge gap in the ssNMR-based bacterial/biofilm research.

Consequently, there is a need to develop and employ robust advanced multidimensional ssNMR techniques for the structural and dynamic characterization of native biofilms.^32^ Here, we aim to address this lack of structural information on bacterial biofilms towards better understanding of molecular structure and organization by using *Pseudomonas fluorescens* Pf0-1 as our model organism. In this work, we utilize conventional ambient-temperature 1D and 2D MAS ssNMR^33^ to successfully detect, characterize, quantify, and better understand the structural details of a native *Pseudomonas* biofilm as a first milestone towards obtaining high-resolution structural as well as qualitative dynamics information. We present novel high-resolution (room-temperature) ssNMR experiments. To our knowledge, this study comprises the first high-resolution 2D ssNMR characterization of an intact *Pseudomonas* colony biofilm without chemical extraction and fractionation of individual biofilm components, and the quantification of the rigid and mobile fractions of the system.

## 2. Methods

### Preparation of biofilm samples for NMR

Details of the *Pseudomonas fluorescens* colony biofilm preparation was described previously.^34-36^ In brief, liquid cultures of *Pseudomonas fluorescens* Pf0-1 were spotted on solid *Pseudomonas* growth agar (PAF) and incubated at 30°C for three days. Colony biofilms were gently scraped off the agar surface and stored at 4°C in microfuge tubes for ssNMR analysis. For the ^13^C detected ssNMR experiments, we utilized thin-walled 3.2 mm rotors. Total amount of biofilm material in the rotor was ∼40-60 mg. We used a benchtop centrifuge to transfer the biofilm material into the rotor, by using simple pipets attached directly onto the rotor and then centrifuged. A large fraction of the native hydrated biofilm material consists of water, and the effective solid material in the rotor is small as determined by a simple wet-biofilm drying test (data not shown). As a result, we analyzed both the wet and dried biofilm samples in fully packed rotors. Drying method increased the amount of total material in the NMR rotor up to ∼10-fold, due to the removal of excess hydration. Drying of the wet biofilm was performed gently at ∼50°C in an oven under atmospheric conditions.

### NMR Spectroscopy

All the MAS ssNMR experiments were performed at 750 MHz Bruker Avance 3 spectrometer equipped with a low-temperature triple-resonance 3.2 mm probe. We employed 10 kHz MAS spinning for all the experiments, and the set temperature was 275 K for all measurements, which corresponds to a sample temperature of around ambient temperature. 3.3 μs and 5 μs pulses were used for ^1^H and ^13^C. For the CP experiments, 1 ms contact time was used with a 70-100% ramp on the proton channel. ∼90 kHz proton dipolar decoupling was applied for all the spectra recorded. The ^1^H chemical shifts were referenced directly to 0 ppm by using DSS as an internal standard added to the NMR samples, and the ^13^C chemical shifts were indirectly referenced by using the ^1^H frequency.^37^

For the wet biofilm samples, the 1D ^13^C INEPT and CP spectra were recorded with 82k and 43k scans to allow for good SNT for analysis by using 1 and 2 seconds of recycle delays, respectively. For the dry biofilm samples, the 1D ^13^C INEPT and CP spectra were recorded with 55k and 8k scans by using 1 second of recycle delay. The S/N ratios were determined by using Topspin 3.6. The signal region was set to 90 to 10 ppm, and the noise region to -70 to -150 ppm for all spectra. The determined S/N values were used to calculate the SNT by dividing S/N by the square root of the time in terms of minutes. The corresponding SNT values are given in Figure 2. The spectral fitting and peak deconvolution were performed by using the ssNake program package.^38^ The 1D INEPT and CP spectra were processed with gaussian broadening window function (with 35 Hz at GB=0.02 in Topspin).

The 2D ^1^H-^13^C INEPT ssNMR spectrum was recorded with 3k and 1.5k transients for wet and dry biofilm samples, respectively. 1 s recycle delay was used and a total of 78 indirect dimension data points were recorded with an increment of 100 μs. The total experiment time was 67 and 33 hours for the wet and dry biofilm samples, respectively. The 2D ^1^H-^13^C CP ssNMR spectrum was recorded with 1k transients for dry biofilm sample in ∼22 hours. 1 s recycle delay was used and a total of 78 indirect dimension data points were recorded with an increment of 100 μs. The solid-state NMR spectra were recorded at 10 kHz MAS and 750 MHz Bruker Avance III NMR spectrometer utilizing a 3.2 mm probe at 275 K. The solution-state 2D ^1^H-^13^C HSQC NMR spectrum of the monomeric soluble FapC was recorded at a 600 MHz Bruker Avance III spectrometer equipped with a 5 mm triple resonance TCI cryoprobe.^39^ 5 mm NMR sample tube was used with a total volume of 550 μl sample and a protein concentration of ∼100 μM at 274 K. The 2D spectrum was processed with gaussian broadening (with 35 Hz at GB=0.025 in Topspin) for direct dimension and by a mixed sine squared for indirect dimension (SSB=3 in Topspin).

## 3. Results

### 3.1. A general solid-state NMR approach to study native bacterial biofilms

Our conventional ssNMR characterization demonstrates the feasibility of detecting sufficient ^13^C NMR signals at natural abundance from the native biofilm samples. **Figure 1A** depicts biofilm formation and the extracellular components that are crucial for maintaining its structural integrity, such as polysaccharides, fibrillar functional amyloid proteins, lipids and eDNA. **Figure 1B** summarizes the two different approaches to record spectra for structural analysis, by conventional room-temperature or low-temperature hyperpolarized DNP-enhanced ssNMR. In this work, we utilized the high-resolution conventional ssNMR approach performed at ambient temperature of ∼300 K to characterize *Pseudomonas* biofilm. This allows the detection of biofilms close to their native *in vivo* conditions. By recording CP or INEPT based ssNMR experiments the rigid and dynamics parts of the biofilm can be quantified and analyzed, respectively, **Figure 1C**. The DNP-enhanced ssNMR spectra on the other hand, result in much larger higher-sensitivity spectra due to hyperpolarization, however, due to the experimental conditions of ∼100 K the mobile fraction of the signals is frozen and only a cumulative rigid spectrum is obtained. Both methods have advantages and disadvantages and could be utilized simultaneously or separately according to the information content needed.

**Figure 1.**
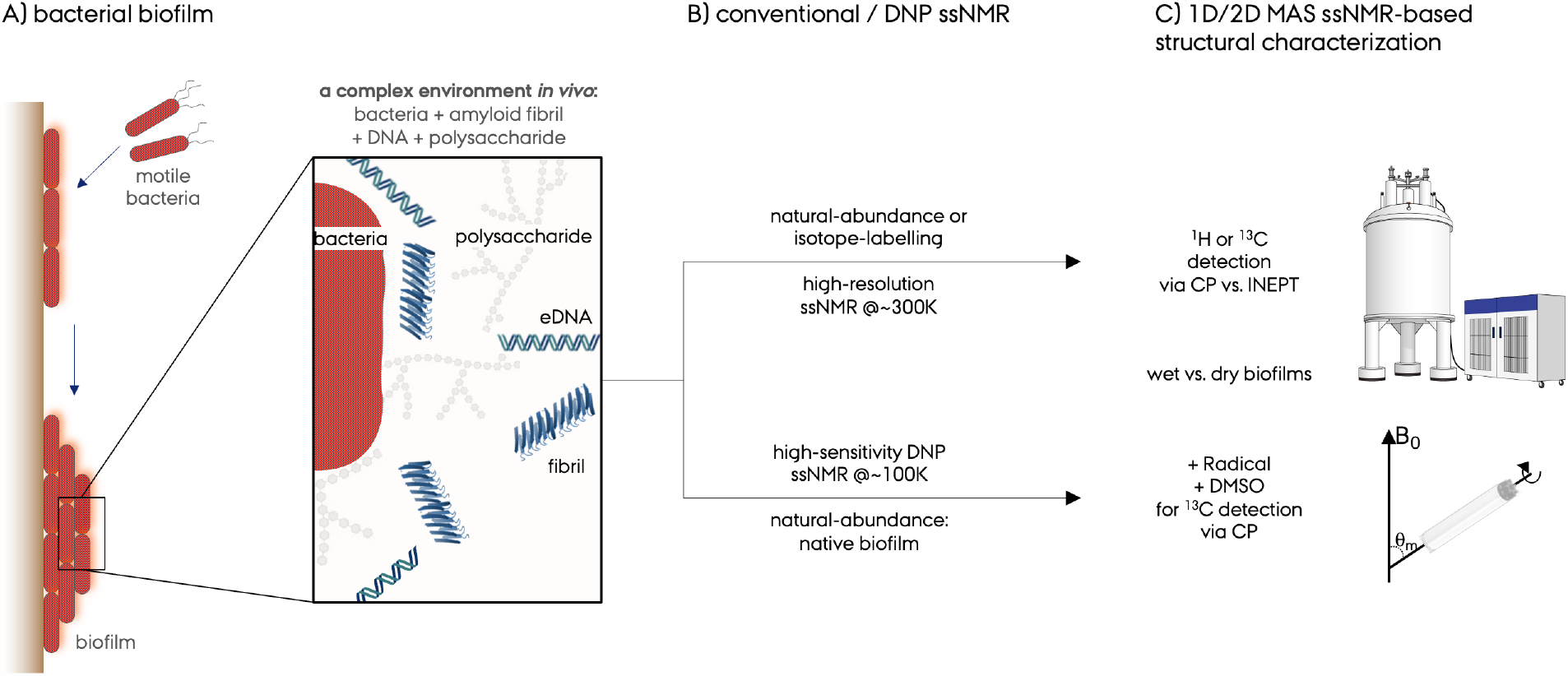
Schematic of the workflow for MAS solid-state NMR study of bacterial biofilms. **A)** A cartoon representation of the biofilm formation, its complex environment, and its components such as bacterial cells, polysaccharides, fibrillar functional amyloid proteins, lipids and extracellular DNA (eDNA). **B)** Two complementary ssNMR approaches that can be utilized to characterize bacterial biofilms, *Pseudomonas* biofilm in this study. The conventional high-resolution ssNMR is usually performed at ∼300 K and the DNP-enhanced high-sensitivity ssNMR is usually performed at ∼100 K. **C)** 1D or 2D ssNMR methods based on CP or INEPT polarization transfer schemes could be applied to record spectra by ^1^H or ^13^C detection depending on the sensitivity, resolution, and isotope-labelling possibilities. The cartoon representation of the NMR spectrometer was obtained from www.wikipedia.com under *nuclear magnetic resonance*.

The ssNMR samples from *Pseudomonas* biofilm were prepared in two different ways, **Figure 2A,B**. First, the native wet biofilms were packed into the ssNMR rotors without any further treatment as close to the *in vivo* condition. In this hydrated biofilm material only a fraction of the total mass is solid due to the access hydration. Second, we gently dried the biofilm at 50ºC to remove the access hydration until a brittle solid is obtained. This drying step improves the sample packing efficiency up to ∼10-fold, and we effectively packed more solid material into the rotor to obtain better signal-to-noise per unit time (SNT). The 1D ^13^C ssNMR spectra recorded with these methods are shown in **Figure 2C,D** at room temperature by CP or INEPT polarization-transfer schemes. Below, 1D and 2D ssNMR results are discussed in detail qualitatively and quantitatively and correlated to biofilm structure and dynamics.

**Figure 2.**
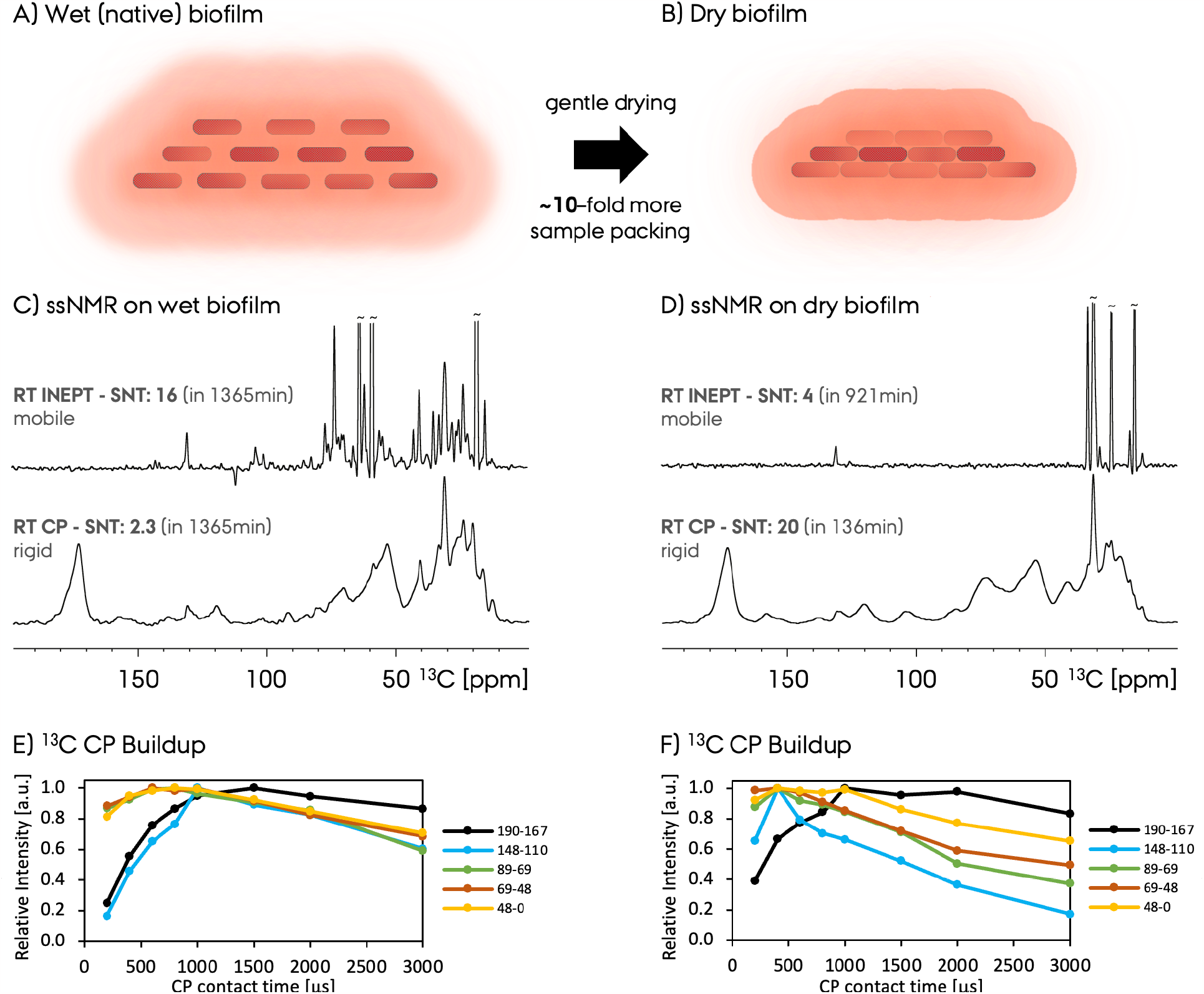
**A**,**B)** Representation of the wet native and dry compact bacterial biofilm. **C**,**D)** 1D ^13^C ssNMR spectra recorded with these methods on wet and dry biofilm preparations. CP and INEPT polarization transfer schemes were used to record the ^13^C ssNMR spectra. The signal to noise ratio per unit time (SNT) values are given along with the total experiment times (determined by (S/N)/(minute)^0.5^ for a direct sensitivity comparison). The CP buildup for different signals in the 1D ^13^C CPMAS spectra (given with the color-coded curves as a functional of contact-time) are shown for **E)** wet and **F)** dry biofilm samples. Each CP buildup curve was normalized to the maximum intensity within that same dataset and different curve intensities are not directly comparable. The five different chemical shift ranges for the integration of signals are given on the side in terms of ppm scale.

### 3.2. 1D ^13^C MAS ssNMR to characterize natural-abundance native *Pseudomonas* biofilm

Conventional room-temperature ^13^C ssNMR spectroscopy on ∼50 mg (in a fully packed 3.2 mm ssNMR rotor) of hydrated native *Pseudomonas* biofilm results in high-sensitivity 1D NMR spectra within ∼1 day (1365 minutes) of total data acquisition, **Fig. 1C**. We recorded the spectra shown in **Figure 2** with ample number of transitions with high SNT to allow quantification more precisely, whereas spectra with sufficient SNT could be recorded in a few hours under these conditions. These spectra were recorded with CP and INEPT which selectively report on the rigid versus mobile/flexible biofilm components, respectively. To our knowledge, this is the first attempt to quantify rigid versus mobile signals/parts of bacterial biofilm, whereas this comparison previously revealed valuable information in a bacterial cell wall study.^31^ These two spectra recorded at room temperature has significantly different sensitivity and resolution. The SNT:16 for the mobile/flexible fraction of the NMR signal recorded by INEPT, whereas the SNT:2.3 for the rigid fraction recorded by CP. Remarkably, a ∼7-fold larger SNT was observed for the mobile spectrum compared to the rigid-fraction. This indicates that under the current experimental conditions the *Pseudomonas* biofilm largely consists of flexible species due to the hydrated nature.

The resolution of the ^13^C INEPT ssNMR spectrum recorded on the wet native biofilm sample contains narrow signals, **Figure 2C top** and **Figure 3B**. The average full-width at half maximum (FWHM) observed for the signals is ∼185 Hz, see **Table 1** for the list of the deconvoluted NMR signals. On the other hand, the ^13^C CP ssNMR spectrum resolution of the same wet biofilm sample is much less with broader and overlapped signals with an average FWHM of ∼490 Hz, **Figure 2C bottom, Figure 3A** and **Table 1**. When the biofilm sample was dried for maximizing ssNMR rotor packing, the ^13^C INEPT ssNMR spectra changes dramatically and only a few signals remain, mostly from the intense lipid signals. Nevertheless, the ^13^C CP spectra are very similar for the wet and dry biofilm samples with the same overall spectral pattern. An increase at the polysaccharide signal intensity at ∼75 ppm is observed for the dry biofilm sample. This indicates increased rigidity of the polysaccharide fraction in the biofilm upon drying, as a result more efficient CP polarization transfer and larger signal. The CP buildup curves for wet and dry biofilm samples are shown in **Figure 2E**,**F**, which further supports changing CP dynamics upon drying as the curves maximizes at shorter contact times except the carbonyl resonance, indicating overall increased rigidity in the sample including protein and carbohydrate fractions. Previously changing of T_1q_ relaxation times, have been correlated to dynamics of carbohydrates in fungal systems,^40^ similar to the current buildup curve changes in ^13^C CP ssNMR experiments. These buildup curves are similar to the behavior obtained on previous polysaccharide and whole cell bacteria work, in which the maximum intensity signal was obtained at ∼500-1000 μs contact time for different sites.^41, 42^

**Table 1:** List of signals determined by deconvolution of the ^13^C MAS ssNMR recorded with **A)** CP or **B)** INEPT, between 13.0 and 181.8 ppm in the spectra shown in Figure 3A,B. Chemical shifts in terms of ppm, percentage integral ratio normalized to the maximum integrals (bold numbers) within the CP or INEPT peaks, and the linewidths are given in the table. A gap in the list indicates that particular chemical shift was not observed in the CP versus INEPT, or vice versa. The chemical shifts of the peaks close to each other are assumed to be from a similar chemical site and listed in the same line in the table. The average FWHM for CP and INEPT spectra are ∼490 and ∼185 Hz, respectively with the processing parameters given in the methods section.

| $^{13}\text{C}$ CP | | | $^{13}\text{C}$ INEPT | | | $^{13}\text{C}$ CP | | | $^{13}\text{C}$ INEPT | | |
| --- | --- | --- | --- | --- | --- | --- | --- | --- | --- | --- | --- |
| ppm | % | FWHM [Hz] | ppm | % | FWHM [Hz] | ppm | % | FWHM [Hz] | ppm | % | FWHM [Hz] |
| 13.0 | 0.181 | 489 | 15.2 | 0.085 | 200 | 84.8 | 0.089 | 697 | 84.6 | 0.003 | 175 |
| 16.6 | 0.445 | 480 |  |  |  |  |  |  | 85.5 | 0.011 | 175 |
| 18.7 | 0.136 | 341 | 18.3 | 0.949 | 150 | 87.6 | 0.010 | 300 | 87.9 | 0.004 | 250 |
| 20.3 | 0.403 | 381 | 20.1 | 0.021 | 300 |  |  |  | 89.8 | 0.002 | 100 |
| 22.0 | 0.731 | 660 | 22.0 | 0.070 | 300 | 92.1 | 0.086 | 482 | 90.7 | 0.002 | 100 |
| 24.2 | 0.425 | 370 | 23.8 | 0.131 | 250 |  |  |  | 93.2 | 0.003 | 125 |
| 26.2 | 0.663 | 500 | 25.6 | 0.082 | 275 | 95.3 | 0.006 | 150 | 95.4 | 0.002 | 100 |
|  |  |  | 26.5 | 0.021 | 100 |  |  |  | 97.5 | 0.006 | 150 |
| 28.4 | 0.426 | 411 | 28.0 | 0.070 | 200 |  |  |  | 98.4 | 0.007 | 150 |
| 31.2 | 0.885 | 396 | 30.8 | 0.264 | 350 | 100.4 | 0.003 | 150 | 101.1 | 0.017 | 175 |
| 33.6 | 0.638 | 579 | 32.9 | 0.067 | 225 | 101.8 | 0.017 | 200 | 102.1 | 0.006 | 225 |
|  |  |  | 35.2 | 0.067 | 178 | 103.6 | 0.014 | 250 | 104.3 | 0.033 | 200 |
| 37.1 | 0.537 | 891 | 37.6 | 0.039 | 400 |  |  |  | 106.1 | 0.008 | 200 |
| 40.7 | 0.146 | 303 | 40.6 | 0.124 | 250 | 116.8 | 0.027 | 332 | 117.3 | 0.005 | 175 |
| 42.0 | 0.510 | 884 | 43.0 | 0.046 | 175 | 119.6 | 0.151 | 617 | 119.9 | 0.004 | 200 |
| 46.1 | 0.189 | 954 | 46.7 | 0.008 | 200 | 122.1 | 0.076 | 878 |  |  |  |
|  |  |  | 47.6 | 0.012 | 200 | 123.8 | 0.013 | 369 |  |  |  |
|  |  |  | 48.6 | 0.011 | 200 | 125.3 | 0.014 | 309 | 125.5 | 0.004 | 150 |
| 50.4 | 0.379 | 800 | 51.1 | 0.015 | 200 | 128.0 | 0.058 | 500 |  |  |  |
|  |  |  | 52.1 | 0.030 | 200 | 128.0 | 0.058 | 500 |  |  |  |
| 53.8 | **1.000** | 800 | 54.9 | 0.047 | 200 | 129.4 | 0.033 | 300 |  |  |  |
| 56.6 | 0.425 | 427 | 56.2 | 0.056 | 200 | 131.0 | 0.085 | 300 | 130.7 | 0.042 | 150 |
| 58.8 | 0.239 | 400 | 58.9 | **1.000** | 150 | 133.1 | 0.021 | 400 | 132.0 | 0.009 | 150 |
| 61.2 | 0.720 | 1050 | 60.9 | 0.011 | 100 | 134.4 | 0.006 | 400 |  |  |  |
|  |  |  | 62.1 | 0.116 | 200 | 136.2 | 0.017 | 300 |  |  |  |
|  |  |  | 64.0 | 0.329 | 150 | 138.6 | 0.043 | 400 |  |  |  |
| 65.9 | 0.405 | 600 | 66.3 | 0.029 | 200 | 141.3 | 0.007 | 200 | 141.4 | 0.006 | 150 |
| 68.1 | 0.055 | 300 | 67.7 | 0.010 | 175 |  |  |  | 143.1 | 0.009 | 150 |
| 70.3 | 0.308 | 600 | 70.0 | 0.036 | 125 |  |  |  | 145.3 | 0.005 | 250 |
|  |  |  | 70.8 | 0.034 | 125 | 153.0 | 0.019 | 400 |  |  |  |
| 72.1 | 0.116 | 615 | 72.0 | 0.051 | 200 | 154.7 | 0.018 | 400 |  |  |  |
| 73.7 | 0.158 | 500 | 73.6 | 0.168 | 175 | 157.4 | 0.043 | 500 |  |  |  |
|  |  |  | 74.5 | 0.005 | 200 | 157.9 | 0.011 | 500 |  |  |  |
| 76.1 | 0.145 | 500 | 75.8 | 0.033 | 250 | 166.1 | 0.010 | 250 |  |  |  |
|  |  |  | 77.1 | 0.061 | 225 | 170.9 | 0.187 | 650 |  |  |  |
| 79.1 | 0.087 | 485 | 78.4 | 0.003 | 150 | 173.3 | 0.720 | 600 |  |  |  |
| 79.9 | 0.025 | 305 | 79.7 | 0.004 | 125 | 176.6 | 0.499 | 800 |  |  |  |
| 81.4 | 0.066 | 340 | 82.8 | 0.013 | 150 | 181.8 | 0.084 | 600 |  |  |  |

**Figure 3.**
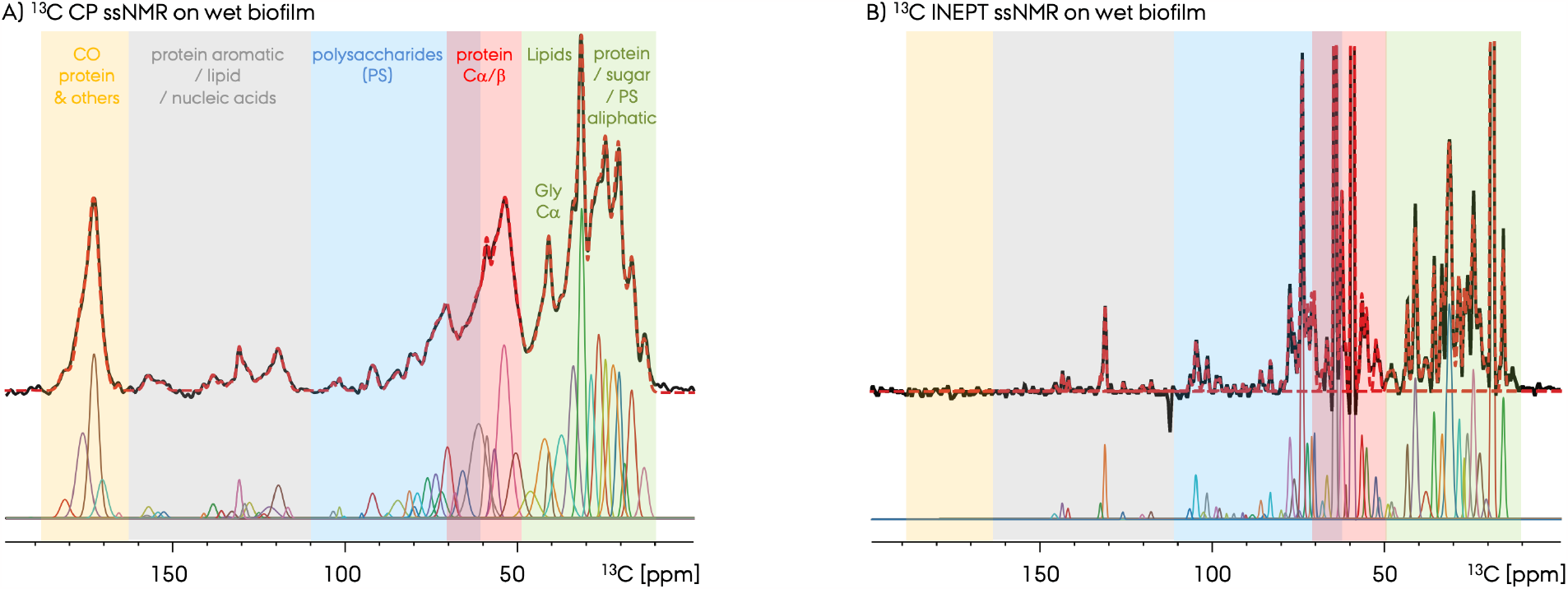
**A)** Quantification of different chemical species in the bacterial biofilm 1D ^13^C MAS ssNMR spectra recorded with CP and INEPT polarization transfer by peak deconvolution. The peaks are determined and fitted by ssNake program. The list of these peaks along with their linewidths and ratios are given below in Table 1. Different tentative group assignments are color coded in the spectra and labeled in A.

**Figure 3 A,B** depicts the deconvolution analysis and tentative group assignments of the carbon chemical shifts observed for the wet native biofilm sample by CP and INEPT ^13^C ssNMR spectra. We utilized ssNake fitting program package,^38^ and was able to fit the spectra to a very good agreement. This allowed us to semi-quantitatively analyze the observed chemical species already in the 1D ^13^C ssNMR spectra and to compare the CP and INEPT spectra. Similar deconvolution of NMR signals approach was shown previously as a valid approach by Schaefer, Cegelski, and coworkers.^43, 44^ We demonstrate here extensive deconvolution and fitting of all peaks in these 1D spectra, as a showcase of extracting detailed information from 1D spectra of complex biological environments. Since the carbon signals from polysaccharides, proteins and other biofilm components have different chemical structure, they are observed at different chemical shift ranges despite several overlapped regions. With this protocol ∼80 unique chemical sites were identified and listed in **Table 1**. The resonance sets identified in the CP and INEPT spectra resemble each other, with still unique differences. The previous studies and growing database on polysaccharide/carbohydrate chemical shifts, help us greatly to tentatively assign the peaks, as well as our recent solution-state NMR assignment of the *Pseudomonas* biofilm forming functional amyloid FapC as a disordered monomer.^39, 43-45^ The 2D ^1^H-^13^C INEPT ssNMR spectrum is the ultimate way to quantitatively separate and analyze these signals, as shown in **Figure 4A**,**B**, and will be discussed below. Nevertheless, this 1D ^13^C spectrum deconvolution approach is still beneficial in particular in combination to the 2D spectrum and is used here to identify chemical species and compare their relative abundance.

**Figure 4.**
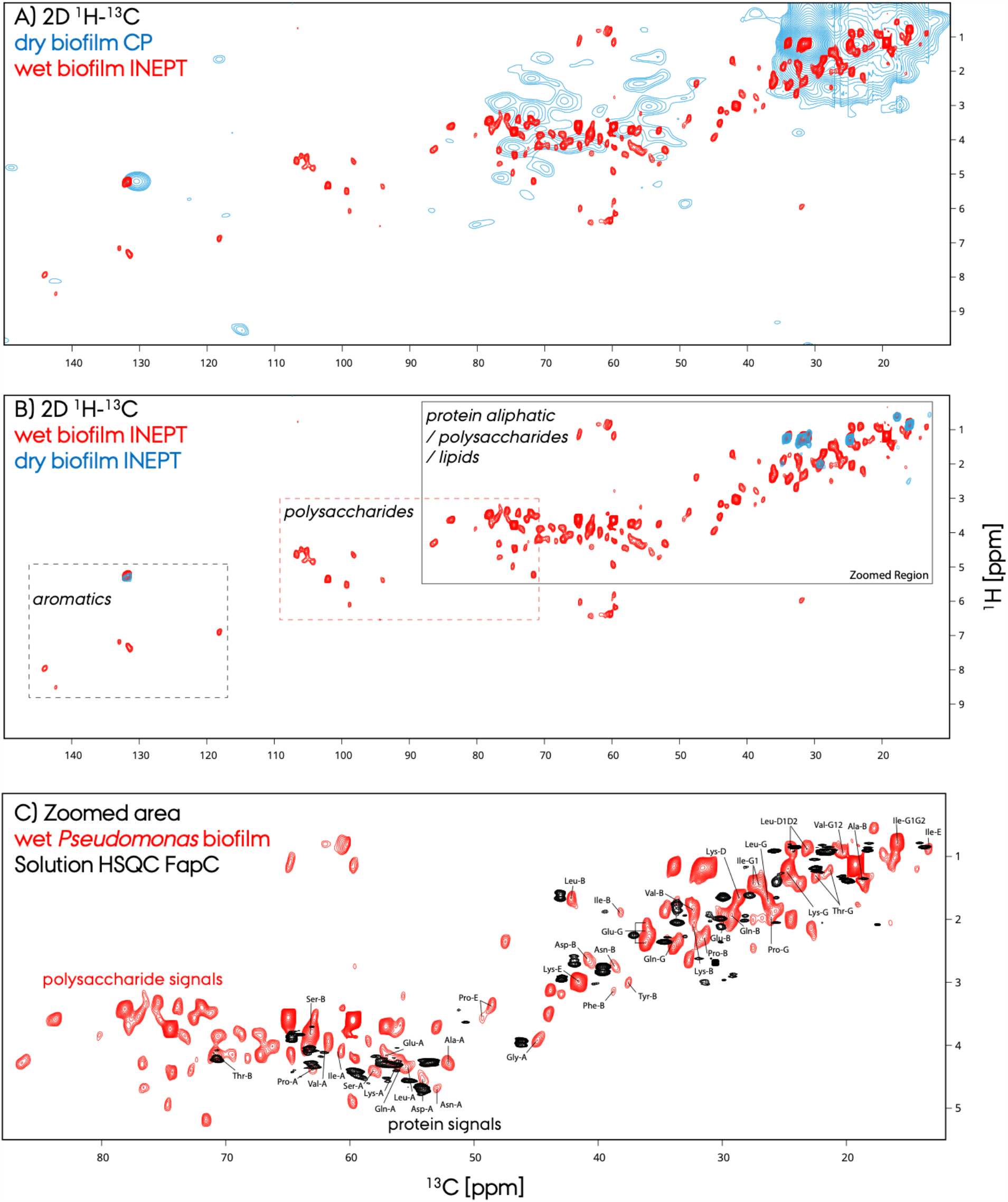
High-resolution 2D ^1^H-^13^C ssNMR correlation spectra for the wet (in red) and dried (in blue) native and natural-abundance *Pseudomonas* biofilm samples recorded with INEPT and CP polarization transfer schemes. **A)** Comparison of the 2D ^1^H-^13^C ssNMR spectra of the dry biofilm recorded with CP (in blue), and of the wet biofilm recorded with INEPT (in red) **B)** The 2D ^1^H-^13^C INEPT ssNMR spectra of wet (in red) and dry (in blue) biofilms. Different signal areas are highlighted in the spectra. **C)** The protein aliphatic / polysaccharide region is shown as a zoom out (in red). The solution-state HSQC NMR spectrum of the biofilm forming functional amyloid FapC as a soluble monomer is overlayed (in black) to indicate the protein signals in the unstructured protein conformations. The protein signals of the biofilm spectra were tentatively assigned by utilizing the FapC spectra and the BMRB database.

The observed chemical shifts are consistent with the polysaccharide, carbohydrate, bacteria, and biofilm studies reported previously.^39, 43-45^ Overall, the carbonyl signals are at ∼190 – 165 ppm, aromatic signals (and/or nucleic acids) from proteins are at ∼165 – 110 ppm, polysaccharides (and/or nucleic acids) are at ∼110 – 65 ppm, protein C_α*/*μ_ are at ∼70 – 50 and glycines C_α_ are at ∼45 ppm, and finally the aliphatic signals from proteins, carbohydrates and lipids are at ∼50-10 ppm. The existence of the carbonyl NMR signal at ∼175 ppm indicates the presence of proteins, as this peak disappears at the INEPT spectrum due to the lack of protons at carbonyl carbons. This is additionally supported by the NMR signals observed for alpha and aliphatic protein carbons at ∼70-10 ppm. Moreover, the C_α_ carbon signals in the 2D ^1^H-^13^C ssNMR spectrum correlate with the correct alpha proton shifts at ∼4.5 ppm, **Figure 4**. This is consistent with the previous 1D NMR studies.^46^ Due to the presence of intense resonance from lipid signals at 18.3, 30.8 ppm and other signals at 58.9 and 64.0 ppm the percentage integral of the other peaks are remarkably small. For the ^13^C CP spectrum the spread of peak integrals is less broad since the spectrum doesn’t have as much of intensity difference as the INEPT spectrum.

### 3.3. 2D ^1^H-^13^C MAS ssNMR spectroscopy for high-resolution native biofilm characterization

The 2D INEPT ^1^H-^13^C MAS ssNMR spectrum shown in **Figure 4A**,**B** is recorded in ∼67 and ∼33 hours for wet and dry biofilms, respectively. Moreover, the 2D CP ^1^H-^13^C MAS ssNMR spectrum shown in **Figure 4A** is recorded in ∼22 hours. The CP-based spectrum comprises signals from the rigid biofilm fractions, whereas INEPT-based spectrum comprises signals from the flexible and mobile wet/dry biofilm species at natural-abundance without isotope-labelling. The spectral sensitivity and resolution of the INEPT-based 2D are excellent with averaged ^13^C resonance linewidths of ∼185 Hz. To our knowledge this spectrum is the first demonstration of multidimensional ssNMR spectroscopy at high-resolution for bacterial biofilms. The CP based 2D spectrum has much less resolution, with an average ^13^C linewidth of ∼490 Hz, which hampers the sensitivity. Different spectral regions comprising signals from protein aliphatic and polysaccharide, polysaccharide and aromatic regions are highlighted in **Figure 4B**. The zoomed out aliphatic region is shown at the bottom for clarity, **Figure 4C**. In **Figure 4B**, the 2D spectra of the wet and dried biofilm samples are shown in red and blue colors, respectively. The dried biofilm sample yielded only a few sharp resonances as also shown in the 1D comparison in **Figure 2**, nevertheless overlapping perfectly with the wet biofilm spectra. To help tentative assigning the protein resonances and differentiate them from the polysaccharide signals for example, we utilized solution-state 2D ^1^H-^13^C HSQC spectrum of the soluble monomeric FapC protein, in black color at the bottom panel. FapC is a biofilm forming functional amyloid from *Pseudomonas* and is an intrinsically disordered protein (IDP) in solution.^39^ The signals from FapC matches the signals of the biofilm sample to a high degree, and we used FapC assignment as the basis of the tentative signal assignment in the biofilm spectrum, along with the BMRB database. In this manner, we accomplished the differentiation of the overlapped regions, where the protein and polysaccharide signals coexist at ∼70 – 10 ppm. The signals observed from the flexible fraction of the biofilm sample (red spectrum) appear to be originating from the less ordered proteins, peptides, or amino acids due to the striking resemblance to the IDP FapC spectrum. There are additional signals in the biofilm spectrum, which we at this point suffice by referring to those as signals from the non-protein fraction of the biofilm. Extracting structural information from the 2D spectrum of the rigid biofilm fraction via CP remains as a challenge due to low resolution.

Taking advantage of the high-resolution 2D ^1^H-^13^C INEPT ssNMR spectra shown in **Figure 4** and utilizing the complex carbohydrate magnetic resonance database (CCMRD), we attempted to determine the types of polysaccharide signals that are observed, **Figure 5**. The *Pseudomonas* biofilm polysaccharides include diverse and complex polymeric substances, including alginate which is widespread across *Pseudomonas* species. The well-studied *P. aeruginosa* strain polysaccharides are composed of Psl, Pel and alginate and others, and are here utilized a guide for signal identification.^8^ In addition, the soluble extracellular polysaccharide composition of our model organism, *P. fluorescens* Pf0-1, was previously quantified.^5, 47^ All in all, by utilizing the ^1^H and ^13^C chemical shifts observed in the 2D INEPT spectrum, we identified several polysaccharide signals from glucose, mannan, galactose, heptose, rhamnan, fucose, and N-acylated mannuronic acid,^8, 25, 48^ which agrees with the previous polysaccharide composition analysis.^47^

**Figure 5.**
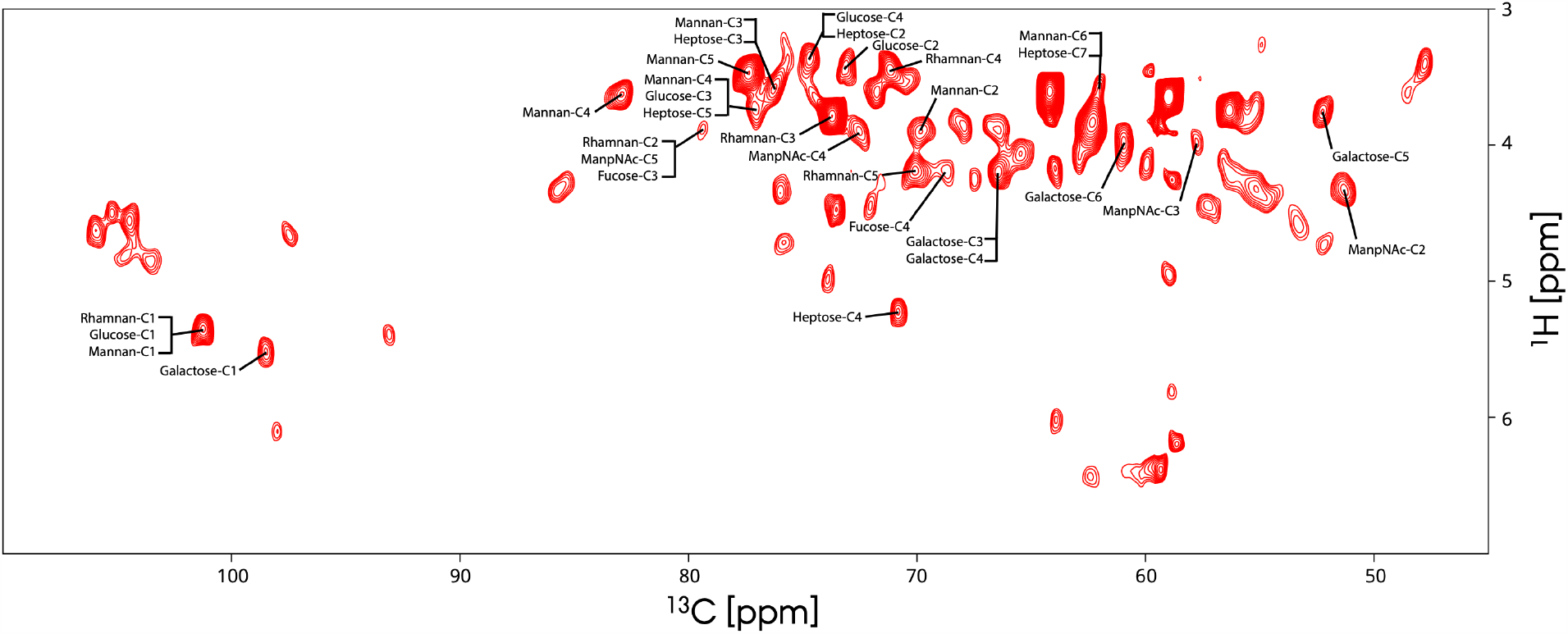
Representation of the tentative polysaccharide ^1^H/^13^C resonance assignments in the INEPT-based 2D ^1^H-^13^C ssNMR spectrum for the wet *Pseudomonas* biofilm sample. We utilized the chemical shift values and the CCMRD database for identification of species. We only judge the presence of a certain polysaccharide species with four to six cross peak pairs identified from a specific candidate. These assignments comprise species such as glucose (CCMRD entry #318), mannan (poly-mannose, #33), galactose (#319), heptose (#327), rhamnan (poly-rhamnose, #329), fucose (CCMRD entry #333), and N-acylated mannuronic acid (ManpNAc #335). To unambiguously assign the resonances and identify the remaining signals, additional experiments are required and will be presented in the future.

## 4. Discussion and Conclusions

We present in this work a high-resolution one- and two-dimensional ssNMR study of the *Pseudomonas fluorescens* biofilm, which is amenable to studying biofilms of any given microbial species including pathologic *P. aeruginosa*. 1D/2D MAS ^13^C-detected ssNMR spectra was utilized to obtain signals from different chemical species in the biofilm and to characterize its molecular composition. This adds valuable structural and dynamics information input to the growing and important research area in structural biology of biofilms, which have been predominantly depended on 1D ssNMR and limit the extend of information that could be extracted from these complex systems.

We propose multidimensional ssNMR spectroscopy as a novel and a powerful tool to study biofilms by selectively obtaining signal from the rigid or mobile fractions of native biofilms. With the presented tentative ssNMR signal assignments, we were able to characterize the bacterial biofilm composition. 1D ssNMR spectra were recorded within several hours to a day with varying SNT and provide insights into the chemical species that exist in the biofilm. The high SNT INEPT based 2D ssNMR spectrum was recorded for the wet biofilm sample within 67 hours and provides resonance identification compared to 1D spectra with more confidence. We anticipate that a 2D INEPT ssNMR spectrum of a lower, but sufficient SNT could be obtained in less than a day, which will provide a fast and high resolution means of detection and differentiation of different carbon pools corresponding to different chemical species. This will open new possibilities for understanding the interaction of proteins and other components (polysaccharides, lipids, and cell-wall) in *Pseudomonas* and other biofilms without the need of isotope labelling or isolating biofilm parts.^28, 46^ We confidently identified several polysaccharide species by analyzing the cross-peaks observed in the 2D spectrum and finding matches in the CCMRD polysaccharide database.

The challenge for future studies to overcome is the sensitivity limitation to characterize the rigid fraction of the biofilm, and similarly, to characterize the dried biofilm with sufficient resolution by multidimensional ssNMR spectroscopy. For the wet biofilm preparation, as shown in **Figure 2**, the sensitivity for the rigid fraction of the biofilm is ∼7 less compared to the mobile fraction. This indicates that a 2D spectrum similar to the one in **Figure 4A**,**B** in terms of sensitivity would need ∼3300 hours (67 hours*(7^2)), which is beyond practicality in experimental time. In addition, since the resolution of the rigid fraction is much less compared to the mobile fraction, **Figure 4A**, the sensitivity is further hampered due to broader line widths at the current moderate MAS frequency conditions. Nevertheless, by using the dry biofilm approach, we increased the amount of biofilm material in the ssNMR rotor and managed to record a CP-based 2D ^1^H-^13^C spectrum, **Figure 4A**. Despite lower resolution obtained, the spectrum overlays well with the INEPT-based 2D. More optimizations of sample preparation and ssNMR spectroscopy are required to utilize the dry biofilm preparations.

DNP enhanced ssNMR would be very beneficial for increasing sensitivity of these experiments to record 2D or even 3D spectra of biofilms.^49, 50^ Such DNP ssNMR approach on fungal systems have been shown successful previously.^16^ Secondly, ultra-fast MAS at >100 kHz would also increase the sensitivity and resolution remarkably and could make it possible to characterize these systems within reasonable time and sufficient resolution. Such a fast MAS approach was shown for bacterial cell wall system.^31^ Finally, a third approach could be isotope-labeling the biofilm systems to increase sensitivity by a factor of ∼100 for ^13^C detection. This method has been used with selectively or fully isotope labeled systems,^42, 43^ however, limits the application of ssNMR to the characterization of lab-grown systems and is not suitable for studying patient derived systems. In summary, we presented a high-resolution ssNMR study to characterize native *Pseudomonas fluorescens* colony biofilm, which could be readily applied to other *in vitro* biofilm model systems and natural biofilms. Wet native and dried compact biofilm sample preparation methods yielded valuable structural insights and up to ∼80/134 distinct chemical sites were identified by peak deconvolution approach in 1D spectra and in the high-resolution 2D spectrum. Based on the CP buildup behavior of the wet and dry biofilm samples, dynamical changes were qualitatively determined. A unique comparison of CP and INEPT ^13^C ssNMR spectra allowed the differentiation of signals from the mobile versus rigid fractions of the biofilm. The high-resolution INEPT based 2D ^1^H-^13^C ssNMR spectrum along with the deconvoluted resonances from the 1D spectra resolved many cross peaks. This allowed the identification of protein signals by utilizing unstructured FapC protein as a template from the polysaccharides signals which appear at different chemical shifts compared to the protein signals. This study represents the first demonstration of high-resolution 2D ssNMR to characterize a native biofilm sample at natural-abundance and without any chemical or physical manipulation and extends the technical capacity of ssNMR in biofilm research. We demonstrate the potential of multidimensional ssNMR in structural studies for identifying chemical sites with much reduced signal overlap. We provide the basis for high-sensitivity and high-resolution structural ssNMR characterization of biofilms and their structurally important components. Our experimental pipeline promises to carry broad impact on related research directions to stimulate new ideas and breakthroughs.

## Data Availability

All data can be requested from the corresponding author.

## Acknowledgements

UA acknowledges financial support from University of Pittsburgh startup funding and the high-field NMR infrastructure at the Structural Biology Department, School of Medicine, University of Pittsburgh. WK was supported by funding from the National Institute of General Medical Sciences of the NIH 1R15GM132856.

## References

(1) Romero, D.; Aguilar, C.; Losick, R.; Kolter, R. Amyloid fibers provide structural integrity to Bacillus subtilis biofilms. Proceedings of the National Academy of Sciences of the United States of America 2010, 107 (5), 2230–2234. DOI: 10.1073/pnas.0910560107.

(2) Akbey, U.; Andreasen, M. Functional amyloids from bacterial biofilms - structural properties and interaction partners. Chemical Science 2022, 13 (22), 6457–6477. DOI: 10.1039/d2sc00645f.

(3) Gebbink, M. F.; Claessen, D.; Bouma, B.; Dijkhuizen, L.; Wosten, H. A. Amyloids--a functional coat for microorganisms. Nat Rev Microbiol 2005, 3 (4), 333–341. DOI: 10.1038/nrmicro1127.

(4) Murray, C. J.; Ikuta, K. S.; Sharara, F.; Swetschinski, L.; Aguilar, G. R.; Gray, A.; Han, C.; Bisignano, C.; Rao, P.; Wool, E. Global burden of bacterial antimicrobial resistance in 2019: a systematic analysis. The Lancet 2022.

(5) Karygianni, L.; Ren, Z.; Koo, H.; Thurnheer, T. Biofilm Matrixome: Extracellular Components in Structured Microbial Communities. Trends in Microbiology 2020, 28 (8), 668–681. DOI: 10.1016/j.tim.2020.03.016.

(6) Bohning, J.; Ghrayeb, M.; Pedebos, C.; Abbas, D. K.; Khalid, S.; Chai, L.; Bharat, T. A. M. Donor-strand exchange drives assembly of the TasA scaffold in Bacillus subtilis biofilms. Nature Communications 2022, 13 (1). DOI: 10.1038/s41467-022-34700-z.

(7) Roske, Y.; Lindemann, F.; Diehl, A.; Cremer, N.; Higman, V. A.; Schlegel, B.; Leidert, M.; Driller, K.; Turgay, K.; Schmieder, P.; et al. TapA acts as specific chaperone in TasA filament formation by strand complementation. Proceedings of the National Academy of Sciences 2023, 120 (17), e2217070120. DOI: 10.1073/pnas.2217070120 (acccessed 2023/07/11).

(8) Flemming, H.-C.; van Hullebusch, E. D.; Neu, T. R.; Nielsen, P. H.; Seviour, T.; Stoodley, P.; Wingender, J.; Wuertz, S. The biofilm matrix: multitasking in a shared space. Nature Reviews Microbiology 2023, 21 (2), 70–86. DOI: 10.1038/s41579-022-00791-0.

(9) Blanco-Romero, E.; Garrido-Sanz, D.; Rivilla, R.; Redondo-Nieto, M.; Martin, M. In Silico Characterization and Phylogenetic Distribution of Extracellular Matrix Components in the Model Rhizobacteria Pseudomonas fluorescens F113 and Other Pseudomonads. Microorganisms 2020, 8 (11). DOI: 10.3390/microorganisms8111740.

(10) Tuttle, M. D.; Comellas, G.; Nieuwkoop, A. J.; Covell, D. J.; Berthold, D. A.; Kloepper, K. D.; Courtney, J. M.; Kim, J. K.; Barclay, A. M.; Kendall, A.; et al. Solid-state NMR structure of a pathogenic fibril of full-length human alpha-synuclein. Nature Structural & Molecular Biology 2016, 23 (5), 409–415. DOI: 10.1038/nsmb.3194.

(11) Asano, S.; Engel, B. D.; Baumeister, W. In Situ Cryo-Electron Tomography: A Post-Reductionist Approach to Structural Biology. Journal of Molecular Biology 2016, 428 (2), 332–343. DOI: 10.1016/j.jmb.2015.09.030.

(12) Plitzko, J. M.; Schuler, B.; Selenko, P. Structural Biology outside the box - inside the cell. Current Opinion in Structural Biology 2017, 46, 110–121. DOI: 10.1016/j.sbi.2017.06.007.

(13) Earl, L. A.; Falconieri, V.; Subramaniam, S. Microbiology catches the cryo-EM bug. Current Opinion in Microbiology 2018, 43, 199–207. DOI: 10.1016/j.mib.2018.02.012.

(14) Narasimhan, S.; Folkers, G. E.; Baldus, M. When Small becomes Too Big: Expanding the Use of In-Cell Solid-State NMR Spectroscopy. Chempluschem 2020, 85 (4), 760–768. DOI: 10.1002/cplu.202000167.

(15) Smith, A. N.; Harrabi, R.; Halbritter, T.; Lee, D.; Aussenac, F.; van der Wel, P. C. A.; Hediger, S.; Sigurdsson, S. T.; De Paepe, G. Fast magic angle spinning for the characterization of milligram quantities of organic and biological solids at natural isotopic abundance by 13C-13C correlation DNP-enhanced NMR. Solid State Nuclear Magnetic Resonance 2023, 123. DOI: 10.1016/j.ssnmr.2022.101850.

(16) Fernando, L. D.; Widanage, M. C. D.; Shekar, S. C.; Mentink-Vigier, F.; Wang, P.; Wi, S.; Wang, T. Solid-state NMR analysis of unlabeled fungal cell walls from Aspergillus and Candida species. Journal of Structural Biology-X 2022, 6. DOI: 10.1016/j.yjsbx.2022.100070.

(17) Chow, W. Y.; De Paepe, G.; Hediger, S. Biomolecular and Biological Applications of Solid-State NMR with Dynamic Nuclear Polarization Enhancement. Chemical Reviews 2022, 122 (10), 9795–9847. DOI: 10.1021/acs.chemrev.1c01043.

(18) Mak-Jurkauskas, M. L.; Bajaj, V. S.; Hornstein, M. K.; Belenky, M.; Griffin, R. G.; Herzfeld, J. Energy transformations early in the bacteriorhodopsin photocycle revealed by DNP-enhanced solid-state NMR. Proceedings of the National Academy of Sciences of the United States of America 2008, 105 (3), 883–888. DOI: 10.1073/pnas.0706156105.

(19) Bajaj, V. S.; Mak-Jurkauskas, M. L.; Belenky, M.; Herzfeld, J.; Griffin, R. G. Functional and shunt states of bacteriorhodopsin resolved by 250 GHz dynamic nuclear polarization-enhanced solid-state NMR. Proceedings of the National Academy of Sciences of the United States of America 2009, 106 (23), 9244–9249. DOI: 10.1073/pnas.0900908106.

(20) Akbey, U.; Franks, W. T.; Linden, A.; Lange, S.; Griffin, R. G.; van Rossum, B. J.; Oschkinat, H. Dynamic Nuclear Polarization of Deuterated Proteins. Angewandte Chemie-International Edition 2010, 49 (42), 7803–7806. DOI: 10.1002/anie.201002044.

(21) Diehl, A.; Roske, Y.; Ball, L.; Chowdhury, A.; Hiller, M.; Moliere, N.; Kramer, R.; Stoppler, D.; Worth, C. L.; Schlegel, B.; et al. Structural changes of TasA in biofilm formation of Bacillus subtilis. Proceedings of the National Academy of Sciences of the United States of America 2018, 115 (13), 3237–3242. DOI: 10.1073/pnas.1718102115.

(22) Jeffries, J.; Thongsomboon, W.; Visser, J. A.; Enriquez, K.; Yager, D.; Cegelski, L. Variation in the ratio of curli and phosphoethanolamine cellulose associated with biofilm architecture and properties. Biopolymers 2021, 112 (1). DOI: 10.1002/bip.23395.

(23) Reichhardt, C.; Joubert, L. M.; Clemons, K. V.; Stevens, D. A.; Cegelski, L. Integration of electron microscopy and solidstate NMR analysis for new views and compositional parameters of Aspergillus fumigatus biofilms. Medical Mycology 2019, 57, S239–S244. DOI: 10.1093/mmy/myy140.

(24) Ghassemi, N.; Poulhazan, A.; Deligey, F.; Mentink-Vigier, F.; Marcotte, I.; Wang, T. Solid-State NMR Investigations of Extracellular Matrixes and Cell Walls of Algae, Bacteria, Fungi, and Plants. Chemical Reviews 2022, 122 (10), 10036–10086. DOI: 10.1021/acs.chemrev.1c00669.

(25) Jennings, L. K.; Dreifus, J. E.; Reichhardt, C.; Storek, K. M.; Secor, P. R.; Wozniak, D. J.; Hisert, K. B.; Parsek, M. R. Pseudomonas aeruginosa aggregates in cystic fibrosis sputum produce exopolysaccharides that likely impede current therapies. Cell Reports 2021, 34 (8). DOI: 10.1016/j.celrep.2021.108782.

(26) Kirui, A.; Zhao, W. C.; Deligey, F.; Yang, H.; Kang, X.; Mentink-Vigier, F.; Wang, T. Carbohydrate-aromatic interface and molecular architecture of lignocellulose. Nature Communications 2022, 13 (1). DOI: 10.1038/s41467-022-28165-3.

(27) Kang, X.; Kirui, A.; Widanage, M. C. D.; Mentink-Vigier, F.; Cosgrove, D. J.; Wang, T. Lignin-polysaccharide interactions in plant secondary cell walls revealed by solid-state NMR. Nature Communications 2019, 10. DOI: 10.1038/s41467-018-08252-0.

(28) Takahashi, H.; Ayala, I.; Bardet, M.; De Paepe, G.; Simorre, J. P.; Hediger, S. Solid-State NMR on Bacterial Cells: Selective Cell Wall Signal Enhancement and Resolution Improvement using Dynamic Nuclear Polarization. Journal of the American Chemical Society 2013, 135 (13), 5105–5110. DOI: 10.1021/ja312501d.

(29) Nygaard, R.; Romaniuk, J. A. H.; Rice, D. M.; Cegelski, L. Spectral Snapshots of Bacterial Cell-Wall Composition and the Influence of Antibiotics by Whole-Cell NMR. Biophysical Journal 2015, 108 (6), 1380–1389. DOI: 10.1016/j.bpj.2015.01.037.

(30) Laguri, C.; Silipo, A.; Martorana, A. M.; Schanda, P.; Marchetti, R.; Polissi, A.; Molinaro, A.; Simorre, J. P. Solid State NMR Studies of Intact Lipopolysaccharide Endotoxin. Acs Chemical Biology 2018, 13 (8), 2106–2113. DOI: 10.1021/acschembio.8b00271.

(31) Bougault, C.; Ayala, I.; Vollmer, W.; Simorre, J. P.; Schanda, P. Studying intact bacterial peptidoglycan by proton-detected NMR spectroscopy at 100 kHz MAS frequency. Journal of Structural Biology 2019, 206 (1), 66–72. DOI: 10.1016/j.jsb.2018.07.009.

(32) Branda, S. S.; Vik, A.; Friedman, L.; Kolter, R. Biofilms: the matrix revisited. Trends in Microbiology 2005, 13 (1), 20–26. DOI: 10.1016/j.tim.2004.11.006.

(33) Chang-Hyeock Byeon, P. C. W., In-Ja L. Byeon, Umit Akbey. Solution-state NMR Assignment and Secondary Structural Propensities of the Full-Length and Minimalistic-Truncated Prefibrillar Monomeric Form of Biofilm-Forming Functional-Amyloid FapC from Pseudomonas aeruginosa. bioRxiv. DOI: 10.1101/2023.01.22.525107.

(34) Evans, A. F.; Wells, M. K.; Denk, J.; Mazza, W.; Santos, R.; Delprince, A.; Kim, W. Spatial Structure Formation by RsmE-Regulated Extracellular Secretions in Pseudomonas fluorescens Pf0-1. Journal of Bacteriology 2022, 204 (10). DOI: 10.1128/jb.00285-22.

(35) Smith, W. P. J.; Davit, Y.; Osborne, J. M.; Kim, W.; Foster, K. R.; Pitt-Francis, J. M. Cell morphology drives spatial patterning in microbial communities. Proceedings of the National Academy of Sciences of the United States of America 2017, 114 (3), E280–E286. DOI: 10.1073/pnas.1613007114.

(36) Kim, W.; Racimo, F.; Schluter, J.; Levy, S. B.; Foster, K. R. Importance of positioning for microbial evolution. Proceedings of the National Academy of Sciences of the United States of America 2014, 111 (16), E1639–E1647. DOI: 10.1073/pnas.1323632111.

(37) Wishart, D. S.; Bigam, C. G.; Holm, A.; Hodges, R. S.; Sykes, B. D. H-1, C-13 AND N-15 RANDOM COIL NMR CHEMICAL-SHIFTS OF THE COMMON AMINO-ACIDS .1. INVESTIGATIONS OF NEAREST-NEIGHBOR EFFECTS. Journal of Biomolecular Nmr 1995, 5 (1), 67–81. DOI: 10.1007/bf00227471.

(38) van Meerten, S. G. J.; Franssen, W. M. J.; Kentgens, A. P. M. ssNake: A cross-platform open-source NMR data processing and fitting application. Journal of Magnetic Resonance 2019, 301, 56–66, Article. DOI: 10.1016/j.jmr.2019.02.006.

(39) Byeon, C. H.; Wang, P. C.; Byeon, I. J. L.; Akbey, U. Solution-state NMR assignment and secondary structure propensity of the full length and minimalistic-truncated prefibrillar monomeric form of biofilm forming functional amyloid FapC from Pseudomonas aeruginosa. Biomolecular Nmr Assignments 2023. DOI: 10.1007/s12104-023-10135-5.

(40) Kang, X.; Kirui, A.; Muszynski, A.; Widanage, M. C. D.; Chen, A.; Azadi, P.; Wang, P.; Mentink-Vigier, F.; Wang, T. Molecular architecture of fungal cell walls revealed by solid-state NMR. Nature Communications 2018, 9. DOI: 10.1038/s41467-018-05199-0.

(41) Reichhardt, C. The Pseudomonas aeruginosa Biofilm Matrix Protein CdrA Has Similarities to Other Fibrillar Adhesin Proteins. Journal of Bacteriology 2023, 205 (5). DOI: 10.1128/jb.00019-23.

(42) Romaniuk, J. A. H.; Cegelski, L. Peptidoglycan and Teichoic Acid Levels and Alterations in Staphylococcus aureus by Cell-Wall and Whole-Cell Nuclear Magnetic Resonance. Biochemistry 2018, 57 (26), 3966–3975. DOI: 10.1021/acs.biochem.8b00495.

(43) Patti, G. J.; Kim, S. J.; Schaefer, J. Characterization of the peptidoglycan of vancomycin-susceptible Enterococcus faecium. Biochemistry 2008, 47 (32), 8378–8385. DOI: 10.1021/bi8008032.

(44) Romaniuk, J. A. H.; Cegelski, L. Bacterial cell wall composition and the influence of antibiotics by cell-wall and whole-cell NMR. Philosophical Transactions of the Royal Society B-Biological Sciences 2015, 370 (1679). DOI: 10.1098/rstb.2015.0024.

(45) Kang, X.; Zhao, W. C.; Widanage, M. D. C.; Kirui, A.; Ozdenvar, U.; Wang, T. CCMRD: a solid-state NMR database for complex carbohydrates. Journal of Biomolecular Nmr 2020, 74 (4-5), 239–245. DOI: 10.1007/s10858-020-00304-2.

(46) McCrate, O. A.; Zhou, X. X.; Reichhardt, C.; Cegelski, L. Sum of the Parts: Composition and Architecture of the Bacterial Extracellular Matrix. Journal of Molecular Biology 2013, 425 (22), 4286–4294. DOI: 10.1016/j.jmb.2013.06.022.

(47) Kim, W.; Levy, S. B.; Foster, K. R. Rapid radiation in bacteria leads to a division of labour. Nature Communications 2016, 7. DOI: 10.1038/ncomms10508.

(48) Jennings, L. K.; Storek, K. M.; Ledvina, H. E.; Coulon, C.; Marmont, L. S.; Sadovskaya, I.; Secor, P. R.; Tseng, B. S.; Scian, M.; Filloux, A.; et al. Pel is a cationic exopolysaccharide that cross-links extracellular DNA in the Pseudomonas aeruginosa biofilm matrix. Proceedings of the National Academy of Sciences of the United States of America 2015, 112 (36), 11353–11358. DOI: 10.1073/pnas.1503058112.

(49) Konig, A.; Scholzel, D.; Uluca, B.; Viennet, T.; Akbey, U.; Heise, H. Hyperpolarized MAS NMR of unfolded and misfolded proteins. Solid State Nuclear Magnetic Resonance 2019, 98, 1–11. DOI: 10.1016/j.ssnmr.2018.12.003.

(50) Akbey, U.; Oschkinat, H. Structural biology applications of solid state MAS DNP NMR. Journal of Magnetic Resonance 2016, 269, 213–224. DOI: 10.1016/j.jmr.2016.04.003.

